# Gamma-interferon-inducible lysosomal thiol reductase maintains cardiac immuno-metabolic homeostasis in heart failure

**DOI:** 10.1101/2022.07.03.498477

**Authors:** Mark Li, Jinxi Wang, Qingwen Qian, Biyi Chen, Zeyuan Zhang, Elizabeth Barroso, Yuan Zhang, Duane D. Hall, E. Dale Abel, Long-Sheng Song, Ling Yang

## Abstract

**Background:** The lysosome is a central player in maintaining immuno-metabolic homeostasis. However, mechanistic insights into the regulation of lysosome-dependent immuno-metabolism in the heart are lacking. Lysosomal reductase Gamma Interferon-Inducible Thiol Reductase (GILT) is the only identified lysosomal reductase that controls diverse sets of lysosomal enzymes and cargoes.

**Methods:** The role of cardiac GILT was assessed by generating a novel genetic mouse model and employing a multidisciplinary approach including surgical interventions, live *in situ* high resolution microscopy, whole-tissue respirometry analysis, unbiased transcriptomic and metabolomic analyses, and various cell biology and biochemical assays.

**Results:** We found that expression and activity of GILT are reduced in hearts from humans and mice with heart failure (HF). Mice with cardiac specific loss of GILT develop late onset systolic HF at baseline. In the setting of nutrient-overload and experimental left ventricular pressure overload conditions, loss of GILT in cardiomyocytes accelerates the development of heart dysfunction. Transcriptomic and metabolic analyses further revealed that cardiac GILT deficiency alters adaptive immuno-metabolic signatures in the heart. Finally, at the cellular level, cardiac GILT deletion impaired mitochondrial respiration, which was in part due to NLR Family Pyrin Domain Containing 3 (NLRP3)-mediated elevation of mitochondrial oxidative stress.

**Conclusions:** Together, these findings identify a causal link between a lysosome-inflammation axis, mitochondrial function and heart failure. Elucidation of these mechanisms will identify novel therapeutic strategies for treating HF.

## INTRODUCTION

Heart failure (HF) is the leading health and economic burden worldwide. Within the U.S. alone, it is estimated that HF will affect more than 8 million Americans by 2030^1,2^. Although available therapies improve symptomatic and clinical outcomes, patients with HF have poor survival prognoses and are subject to recurrent hospitalizations^3^. Therefore, there is an urgent need to identify critical molecular regulators that could be targeted to promote cardioprotection to ultimately improve treatment and management options for HF patients.

Although heterogeneous in etiology and diagnostic criteria, HF is characterized by abnormalities of cardiac structure and/or function, including pathological structural remodeling (hypertrophy and dilatation), impaired cardiac electrophysiological conductance, and pulmonary congestion ^4^. The cellular signaling events that lead to these pathological alterations have been exclusively investigated. Recently, emerging evidence suggested that dysregulation of mitochondrial metabolism and immune responses are two critical factors involved in HF pathogenesis^5–8^. The heart is an ever-demanding energy organ comprised of high mitochondrial content, which produces the enormous amounts of ATP needed to maintain contractile function^6,7^. As such, impaired energy production and aberrant metabolic reprogramming contribute to cardiac dysfunction and progression of HF in humans and animal models^6,8,9^. Notably, targeting mitochondria has been proposed as a potential option for treating HF in human patients^6,9,10^. Indeed, in two randomized double-blind clinical trials, either testing the efficacy of Coenzyme Q_10_, an electron carrier^11^, or elamipretide, a mitochondria-targeted tetrapeptide^12^, have shown the potential for improving systemic and cardiovascular parameters in patients with HFrEF. In addition, cardiac injury triggers immune responses and physiological inflammation to promote tissue and cellular repair^8,13,14^. However, compromised immune modulation promotes aberrant inflammation, further damaging the heart and accelerating HF progression. Supporting this notion, a randomized double-blind trial of a monoclonal antibody against IL-1β, Canakinumab, showed reduced recurrent cardiovascular events in patients with previous myocardial infarction compared to placebo-treated patients^15^. However, despite the understanding that disruption of mitochondrial metabolism and immune responses are central features that control the progression of HF^8^, little is known about the cardiac immune-metabolic interplay and its functional significance in the setting of HF.

Intracellularly, the lysosome fine-tunes organelle quality control, nutrient sensing, membrane repair and macromolecule catabolism^16–20^. In parenchymal cells, disruption of lysosome-mediated metabolic homeostasis has been described in diverse pathologies. For example, we and others have shown that defective lysosomal function contributes to obesity-associated hepatic insulin resistance^21–23^. In contrast, improving lysosomal function ameliorates steatosis and improves glucose homeostasis in livers of obese mice ^21,24^. The lysosome also serves as a central hub for innate and adaptive immunity^25,26^. For example, in the context of tissue injury, leukocytic lysosomes play key roles in controlling pathogen elimination, antigen presentation, inflammasome activation, and inflammatory lipid metabolism^25–29^. Regarding inflammation, lysosomal dysfunction triggers adipose inflammation^30^ and lysosome-mediated toll-like receptor 4 (TLR4) degradation prevents non-alcoholic steatohepatitis (NASH) in both mice and monkeys^31^. However, in the context of HF, it is unknown whether and how defective cardiac lysosomes drive the immuno-metabolic imbalance.

Lysosomal dysfunction has been implicated in a wide range of cardiovascular diseases (CVDs)^32,33^. However, delineating the physiological and pathological roles of lysosomes *in vivo* has been challenging. This is in part due to the complexity of the organelle, which contains more than 60 enzymes that act upon a multitude of substrates^19,34^. As such, studies of individual lysosomal enzymes using gain- or loss-of-function in mice have resulted in contradictory observations regarding cardiac outcomes under stress conditions^35–43^. Gamma-Interferon-Inducible Lysosomal Thiol Reductase (GILT) holds a central place in maintaining lysosomal homeostasis, as it is the only identified lysosomal reductase that controls activities of a diverse set of lysosomal enzymes^44,45^. To date, GILT has been primarily recognized as a regulator of immune cell antigen presentation ^44^, but its functional role in cardiomyocytes in the context of HF is essentially unknown. Notably, a single nucleotide polymorphism (SNP) in the coding sequence of *IFI30*, which encodes GILT, is associated with increased CVD risk factors^46^. Here we report a protective role of cardiac GILT-mediated lysosomal function in maintaining immuno-metabolic homeostasis in HF.

## MATERIALS AND METHODS

### Human samples, mouse models and animal experiments

The human heart samples were obtained from the UPenn Biobank. Animal experiments were carried out in accordance with the Guide for the Care and Use of Laboratory Animals (National Institutes of Health publication 85–23, revised 1996) and were approved by the Institutional Animal Care and Use Committee at the University of Iowa. The *Ifi30^fl/fl^* mice were generated by Biocytogen (Wakefield, MA), and achieved by targeting the endogenous *Ifi30* locus with the CRISPR-Cas9 technology. LoxP elements were inserted upstream of exon 2 and downstream of exon 7, and Cre-mediated recombination results in a null protein. Following the Cas9/sgRNA and targeting vector construction, the zygote microinjection was carried out, the founder was genotyped, and the mice were back-crossed with C57BL/6J genetic background mice for >5 generations. Homozygous *Ifi30* mutants were crossed with Myosin-heavy-chain-(MHC)-Cre mice to induce cardiomyocyte-restricted deletion of *Ifi30*. Littermate mice were used as controls in the outlined experiments.

Transverse aortic constriction (TAC) was induced in nine to ten-week-old male C57BL/6J mice, as described previously^98^. Briefly, mice were anesthetized with a ketamine (100 mg/kg) and xylazine (5 mg/kg) mix by intraperitoneal injection. Aortic constriction was performed by tying a 7–0 nylon suture ligature against a 27-gauge needle. For the sham group, the aortic arch was visualized but not banded.

### Echocardiography

Transthoracic echocardiograms were performed on conscious mice in the University of Iowa Cardiology Animal Phenotyping Core Laboratory, using a Vevo 2100 Imager (Visual Sonics), as described previously^99,100^. The echocardiographer was blinded to the treatments and genetic backgrounds of the animals. 2D imaging and M-mode echocardiography were performed in the left ventricle (LV) short- and long-axis planes. Left ventricular end-diastolic volume (LVEDV), end-systolic volume (LVESV), and ejection fractions (LVEF) were calculated with the area-length method.

### Isolation of primary cardiomyocytes

Mouse hearts were mounted on the Langendorff apparatus and single ventricular myocytes were isolated by using the collagenase type II (Worthington Biochem.) and protease method described previously ^101^. Ventricular myocytes were placed on glass coverslips coated with laminin in plating medium containing the following: Minimum essential media (ThermoFisher), Insulin-transferrin-selenium (1x, ThermoFisher, 51500056,), penicillin/streptomycin (1x, ThermoFisher, 15140122), L-Glutamine (2 mM, ThermoFisher, 25030081), Beta-mercaptoethanol (Fisher scientific, 44-420-3250ML), NaHCO_3_ (Sigma, 36486), BSA (Sigma, A7030), Blebbistatin (0.832 μM, Sigma, B0560), FBS (10%, Fisher scientific, 16-000-044) and HEPES (Fisher scientific, 15-630-106). Cells were incubated at 37°C supplemented with 5% CO_2_. Three hours after seeding, plating medium was replaced with culture medium (plating medium without FBS).

### Quantitative RT-PCR (qRT-PCR)

Total RNA was isolated using the Trizol reagent (Invitrogen, 15596026, Carlsbad, CA, USA) and reverse transcribed into cDNA using a PrimeScript RT Master Mix (Takara RR036A, Kusatsu, Shiga, Japan). Quantitative real-time RT-PCR analysis was performed using TB Green Premix Ex Taq (Clontech, RR420A, Mountain View, CA, USA). The following diluted primers were used: *HPRT*-F: 5’-CAT TAT GCT GAG GAT TTG GAA AGG-3’; *HPRT*-R: 5’-CTT GAG CAC ACA GAG GGC TAC A-3’; *IFI30*-F: 5’-TCC AAT GCA CCG CTT GTC AAT-3’; *IFI30*-R: 5’-ACC TTG TTG AAT TTG CAC TCC TC-3’; *Hprt*-F: 5’-CAGTCCCAGCGTCGTGATTA-3’; *Hprt*-R: 5’-GGCCTCCCATCTCCTTCATG-3’; *Ifi30*-F: 5’-GCT TGT CGC TAC TTC CTC GT-3’; *Ifi30*-R: 5’-ATG GTT AGG AAC GCT GCC TC-3’. For mouse tissues, standard curve was generated from pooled samples and included in every reaction. For the human heart tissue, ΔΔCt analysis was used.

### Western blot analysis

Heart tissue and cells were mechanically homogenized in a lysis buffer (50 mM Tris-HCl pH 7.5, 2 mM EDTA, 5 mM EGTA, 30 mM NaF, 40 mM β-glycerophosphate, 10 mM sodium pyrophosphate, 0.3% NP-40) and freshly added protease inhibitor cocktail (Sigma-Aldrich, P8340). Protein concentration was determined by Pierce BCA kit (Thermo Fisher Scientific, 23225) and adjusted samples were subjected to SDS-polyacrylamide gel electrophoresis followed by an overnight transfer onto PVDF membranes (Bio-Rad Laboratories, 1620177). Membranes were immunoblotted with antibodies overnight at 4°C as follows: NLRP3 (Adipogen AG-20B-0014-C100; 1:500), Caspase-1 (Adipogen AG-20B-0042-C100; 1:1,000), OxPhos rodent antibody cocktail (ThermoFisher 1:5,000). Secondary antibodies were horseradish peroxidase-conjugated goat anti-mouse-IgG (Santa Cruz Biotechnology, sc-2005) and horseradish peroxidase-conjugated anti-rabbit-IgG (CST 7074) at 1:5,000 dilution for 1 hour at RT. The signal was detected using the ChemiDoc Touch Imaging System (Bio-Rad Laboratories), and densitometry analyses of western blot images were carried out with the Image Lab software (Bio-Rad Laboratories).

### HexB and GILT enzyme activity assay

The HexB enzyme activity assay was carried out as before^22^, and reactions without the substrate were used as a negative control. GILT enzyme activity assay was modified from a protocol previously described^48^. The non-ionic lysis buffer was adjusted to pH 4.5 and heart tissue was homogenized mechanically on ice. 300 ng of F(ab’)2-Rabbit anti-Rabbit IgG (H+L) (CST, 4412S, Lane Danvers, MA, USA) was diluted in 0.1% Triton X-100 (Sigma-Aldrich, T8787, St. Louis, MO, USA) and incubated at 95°C for 5 min. 15 μg of lysate was pre-incubated with 25 μM DTT (Sigma-Aldrich, 43816, St. Louis, MO, USA) at 37°C for 10 min. The boiled antibody was mixed with the pre-activated lysate and the reaction was terminated after 1 hr with 5 mM iodoacetamide (Sigma-Aldrich, I1149, St. Louis, MO, USA). The reaction was resolved on SDS-PAGE with non-reducing conditions. Images were acquired with Bio-Rad GS-250 Molecular imager and analyzed using the Image Lab software (BioRad, Hercules, CA, USA).

### IL-1β secretion in primary cardiomyocytes

Isolated primary cardiomyocytes were incubated in culture medium (0.25 mL per reaction) for 30 min at 37°C prior to measuring constitutive secretion of IL-1β. Cells were then treated with lipopolysaccharide (500 ng/mL, Sigma, O55:B5 L2880) for 50 min followed by a 10 min incubation with ATP (5 mM, RPI, A30030-1.0). Conditioned medium was collected and IL-1β levels were assessed with ELISA MAX^TM^ Deluxe Set Mouse IL-1β (Biolegend, 432604) following manufacturer’s instructions.

### Mitochondrial ROS imaging in primary cardiomyocytes

Isolated primary cardiomyocytes were seeded on 22 mm round coverslips (ThermoFisher Scientific, 174977) pre-coated with laminin for 24 hr prior to treatment with MCC950 (1 μM, 2 hr, Invivogen, inh-mcc) followed by 15 min exposure to isoproterenol (100 nM). Cells were incubated with MitoSOX^TM^ (50 nM, ThermoFisher Scientific, M36008), fixed with 4% paraformaldehyde and counterstained with DAPI before mounting and imaging with a confocal microscope (Zeiss LSM 700, Carl Zeiss MicroImaging Inc., Germany). Images of single z stacks were obtained and analyzed with ImageJ software (NIH, Rockville, MD) and total intensity was normalized to the cell surface.

### High-resolution respirometry

High-resolution oxygen consumption was carried using the OROBOROS Oxygraph-2k (O2k, Oroboros instruments, Innsbruck, Austria). Freshly resected left ventricular tissue (~3-5 mg per sample) was pulled into fibers and permeabilized in the presence of saponin (50 μg/mL, Sigma, 47036) 30 min at 4°C in BIOPS buffer containing the following: 7.23 mM K2EGTA, 2.77 mM CaK2EGTA, 20 mM imidazole, 0.5 mM DTT, 20 mM Taurine, 5.7 mM ATP, 14.3 mM PCr, 6.56 mM MgCl_2_-6H_2_O, 50 mM MES, pH adjusted to 7.1 with 5 N KOH (reagents purchased from Sigma). The digested fibers were then incubated for 10 min at 4°C in buffer Z’ containing the following: 105 mM K-MES, 30 mM KCl, 10 mM KH_2_PO_4_, 5 mM MgCl_2_-6H2O, 2.5 mg/mL BSA-FA-free, pH adjusted to 7.4, and 1 mM EGTA was added prior to the experiment (reagents were purchased from Sigma). Measurements were recorded in the presence of 2.5 mM ADP (state 3 respiration, Sigma, A2754) and the absence of ADP (state 2 respiration) while providing palmitoyl-carnitine (0.075 mM, Sigma, 61251) and malate (1 mM, Sigma, 1613881). MCC950 (5 μM) or MitoTEMPO (50 μM, Sigma, SML0737) were added to the fibers during the saponin incubation as well as the subsequent incubation in buffer Z’ (total of 1 hour).

### *In situ* calcium imaging and in situ T-tubule imaging

Calcium imaging in the heart was carried out as previously described^102^. Briefly, excised hearts were perfused with Rhod-2 AM (0.3 mM, Bioquest, 21060) containing Kreb-Henseleit’s (KH) solution (in mM: 120 NaCl, 24 NaHCO_3_, 11.1 glucose, 5.4 KCl, 1.8 CaCl_2_, 1 MgCl_2_, 0.42 NaH_2_PO_4_, 10 taurine, 5 creatine, oxygenated with 95% O_2_ and 5% CO_2_) at room temperature for 30 min via retrograde Langendorff perfusion system^103^. Hearts were later transferred to another Langendorff apparatus (37 °C) attached to the confocal microscope system after Rhod-2 loading was completed. The heart was placed onto a recording chamber for in situ confocal imaging (line-scan) of Ca^2+^ signals from epicardial myocytes under sinus rhythm. To avoid motion artifacts during Ca^2+^ imaging, Blebbistatin (10 μM, Sigma, B0560) and BDM (2, 3-butandion-monoxim, 10 mM, Sigma) were added to the perfusion solution. The confocal line-scan images were acquired at a rate of 1.93 msec per line (LSM510, Carl Zeiss MicroImaging Inc., Germany). Ca^2+^ transients were autonomously elicited by electrical signals from sinus atrial node and with field stimulation at various frequencies.

*In situ* imaging of T-tubules was carried out as previously described^56^. T-tubules from intact hearts were stained with MM 4-64 (2.5 μM, AAT BioQuest) in Ca^2+^ free Tyrode solution via Langendorff perfusion at room temperature for 30 min as described previously (*53, 54*). The structure of T-tubules was visualized with a confocal microscope (LSM510, Carl Zeiss MicroImaging Inc., Germany) using a 63× lens. Quantitative analysis of T-tubule integrity was processed with the IDL image analysis program as previously reported. The power value (TTpower) reflects the strength of the regularity of T-tubule organization.

### Immunohistochemistry in the left ventricle

Hearts were quickly excised and embedded in Tissue-Tek® O.C.T. compound (Sakura Finetek USA, 4583), which was then frozen in 2-Methylbutane (Sigma, M32631) over liquid nitrogen. Hearts were sectioned into ~8-10 μm slices and fixed in 4% paraformaldehyde for 15 min at room temperature. Next, samples were washed with PBS, blocked in PBS containing CaCl_2_, Triton X-100 (Sigma, X114), and goat serum (Thermo Fisher, 31872) for 1 hour. Primary antibodies were diluted in blocking buffer at 1:100-1:300 and incubated overnight at 4°C in a humidified chamber. Secondary antibody conjugated to Alexa 488/568 (Thermo Fisher, A32731, A-11011) was prepared in blocking buffer at 1:300 and added to the sample for 1 hour at room temperature. Images were acquired with a confocal microscope (Zeiss LSM 700/880, Carl Zeiss MicroImaging Inc., Germany).

### Masson Trichrome staining

Hearts were embedded in OCT and frozen with precooled 2-Methybutane (Sigma, M32631). Heart sections (8 μm) were stained with hematoxylin and eosin (H&E) and Masson’s Trichrome staining using standard procedures. Fibrosis was quantified as a fraction of the total cardiac area using ImageJ software (NIH, Rockville, MD).

### GC-MS steady state metabolomics in the heart

Data were acquired from an Agilent 6545 UHD QTOF interfaced with an Agilent 1290 UHPLC. Metabolites were separated by using a Millipore Sigma SeQuant ZIC-pHILIC (150 mm x 2.1 mm, 5 μm) column. Solvents were A, 95% water in acetonitrile with 10 mM ammonium acetate and 5 μM phosphate, and B, 100% acetonitrile. A flow rate of 200 μL/min was applied with the following gradient (minutes, %B): 0, 94.7%; 2, 94.7%; 27, 36.8%; 35, 20.0%; 37, 20.0%; 39, 36.8%. For all experiments, 2 μL of metabolic extract was injected. MS parameters were as follows: gas, 200°C 4 L/min; nebulizer, 44 psi; sheath gas, 300°C 12 L/min; capillary, 3kV; fragmentor, 100V; scan rate, one scan/s. MS detection was carried out in both positive and negative modes with a mass range of 65–1,700 Da. Identifications were established by comparing the retention times and fragmentation data of compounds to model standards. All raw data files were converted into mzXML files by using msconvert. Data analysis was performed by using either Agilent’s Profinder or in-house R packages.

### Isolation of RNA and data analysis of bulk RNA sequencing

For each experimental group, total RNA was extracted from fresh hearts from 3 individual mice using Trizol and cleaned up with a RNeasy Mini Kit^23^. Quality control of isolated RNA was determined by DropSense 16 (Trinean). RNA sequencing was performed by Iowa Institute of Human Genetics (University of Iowa) using Illumina Novaseq6000 system. RNA-seq reads were quality checked using the FastQC tool (version, 0.1.2) and trimmed using the Java software package Trim Galore! (Version 0.6.5, Babraham Bioinformatics, Babraham Institute, Cambridge, UK). The reads were mapped to the Ensembl mouse (mm10/GRCm38) genomes using STAR software (version 2.7.10)^104^. Transcripts abundance was quantified with Salmon (version 1.8)^105^. The R Bioconductor package tximport were used to collapse abundance estimates from the transcript level to the gene-level (version 1.22)^106^. Differentially expressed genes (DEGs) were identified by applying the R/Bioconductor package Deseq2 (version 1.34.0)^107^. Enriched pathways represented by the DEGs were identified by gene set enrichment analysis (GSEA, version 4.1.0) and Enrichr^108^. The difference of gene between groups was based on adjusted P value <0.05.

### Statistics

Data were analyzed in GraphPad Prism version 9.3.1. Data are presented as mean ± standard error of the mean (SEM). The sample size was chosen based on the lab’s previous experience and other reports in the field. To ensure normality, the Shapiro-Wilk test was used in RStudio for each dataset, wherein the p-value > 0.05 dictated normally distributed datasets. Following the Shapiro-Wilk test, a parametric test was carried out. Multiple group comparisons were calculated by 2-way ANOVA with Tukey post-hoc test. Pairwise testing was carried out by the two-sided unpaired Student’s t-test.

## RESULTS

### GILT is dysregulated in HF in humans and mice

To evaluate the pathophysiological relevance of GILT in the setting of HF, we first measured the transcript level of *IFI30* in heart tissues from human donors with HF with reduced ejection fraction (HFrEF; EF<30%) and donors without HF. As shown in Fig. 1A, *IFI30* was significantly reduced in human failing hearts. We further performed an *in silico* analysis^47^ and predicted deleterious effects of an R-Q SNP associated with CVD on GILT’s enzymatic activity (Sfig. 1A). To determine the impact of cardiac stress on the function of GILT, we measured *in vitro* thiol reductase activity in heart tissue from mice with advanced HfrEF (EF<30%) using fluorophore-labeled-F(ab′)_2_ as a substrate at pH 4.5^48^. As shown in Fig. 1B&C, GILT enzymatic activity was diminished in mouse hearts with HfrEF. These data indicate that advanced HF is associated with GILT deficiency in the heart. To determine the functional impact of GILT deficiency in the heart, we next assessed cardiac function in mice with whole body GILT deletion (GILT^−/−^) by echocardiography. As shown in Fig. 1D-F, mice with germline deletion of GILT exhibited significantly increased heart mass and reduced EF in lean male mice, indicating a potential role for GILT in maintaining systolic function. Taken together, these results demonstrate that GILT is necessary for normal heart function.

**Figure 1.**
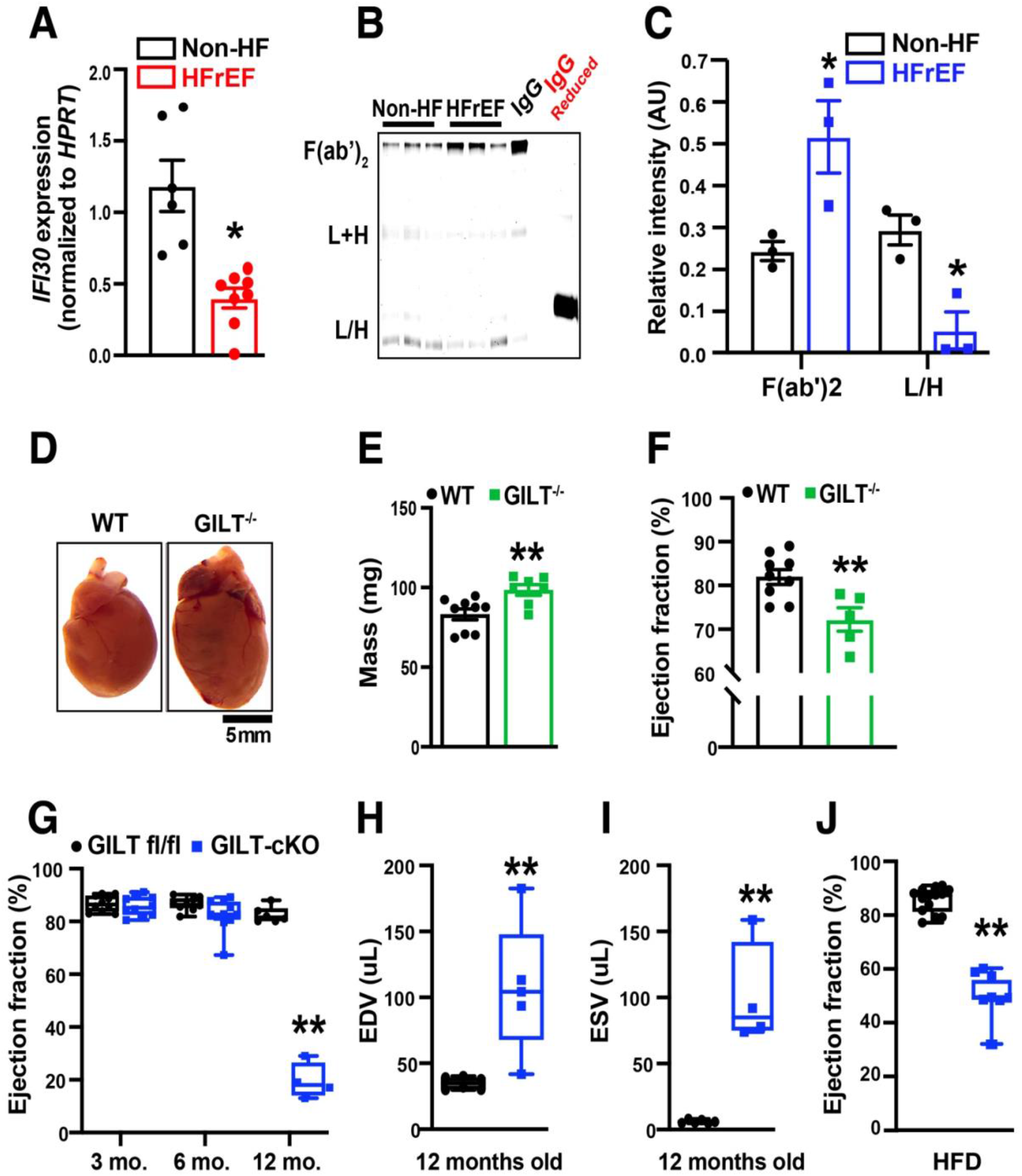
GILT is required for maintaining cardiac function in HF. **A**. Expression of the *IFI30* transcript in hearts from human subjects without heart failure (Non-HF) and with HF with reduced ejection fraction (HFrEF, EF of <30%) measured by qRT-PCR. Data were normalized to *HPRT*; n= 6-8 subjects/group. **B**. Enzyme activity of GILT in cardiac lysates from Non-HF and HFrEF (EF<30%). F(ab’)_2_ and its reduced forms (L+H & L/H) were detected after incubating the substrate with the lysate under acidic conditions; IgG indicates F(ab’)_2_ alone under non-reducing and reducing conditions; each lane represents a reaction from an individual mouse heart. **C**. Quantification of B. **D&E**. Representative images and wet mass of hearts from wild-type (WT) or germline deletion of GILT (GILT^−/−^); n= 6 aged-matched GILT^−/−^ and littermate control (WT) male mice. **F**. Ejection fraction (EF) in WT and GILT^−/−^ mice on regular chow measured by echocardiography; n = 6 aged-matched littermate male mice. **G**. EF in GILT fl/fl and GILT-cKO male mice at indicated ages; n = 4-11 age-matched male littermate mice per group. mo: months. **H**. End-diastolic volume (EDV) and **I**. End-systolic volume (ESV) measured by echocardiography in GILT fl/fl and GILT-cKO male mice at 12 months of age; n = 5-6 age-matched male littermate mice per group. **J**. EF measured by echocardiography in GILT fl/fl and GILT-cKO male mice fed a HFD for 24 weeks; n= 8-14 age-matched male littermate mice per group.Data are presented as means ± SEM. * Indicates genetic effect or HF effect in (A). Two-way ANOVA with Tukey’s multiple comparisons test was utilized for panels (C) and (G), Student’s t-test was utilized for the rest of the panels. * Indicates p-value < 0.05, ** indicates p-value < 0.01.

### Cardiac GILT is required for maintaining the structure and function of the heart

HF progression involves cellular reprogramming in different cell types within the heart in addition to dynamic interplays across tissues and organs^13,49^. To evaluate the necessity of cardiomyocyte GILT in HF pathophysiology, we generated a mouse strain with loxP sites flanking exons 4 through 7 of the *Ifi30* gene (GILT fl/fl; C57BL/6J background). We then deleted *Ifi30* in the heart (GILT-cKO) by crossing GILT fl/fl mice with α-Myosin heavy chain (α-MHC) Cre mice (Sfig. 1B&C). Young male and female GILT-cKO mice (3 months old) displayed normal cardiac systolic function by echocardiography (Fig. 1G, and Sfig. 2A&B). It is well recognized that age is a risk factor for HF^50^. Indeed, we observed that in male mice at 12 months of age, loss of GILT in cardiomyocytes significantly decreased heart function relative to GILT fl/fl mice (Fig. 1G-I and Sfig. 2B). Obesity also represents a significant risk factor for HF development^8^. Indeed, we found that diet-induced obesity (DIO) promoted the development of HF in male GILT-cKO mice compared to their littermate controls (Fig. 1J, SFig. 2C).

**Figure 2.**
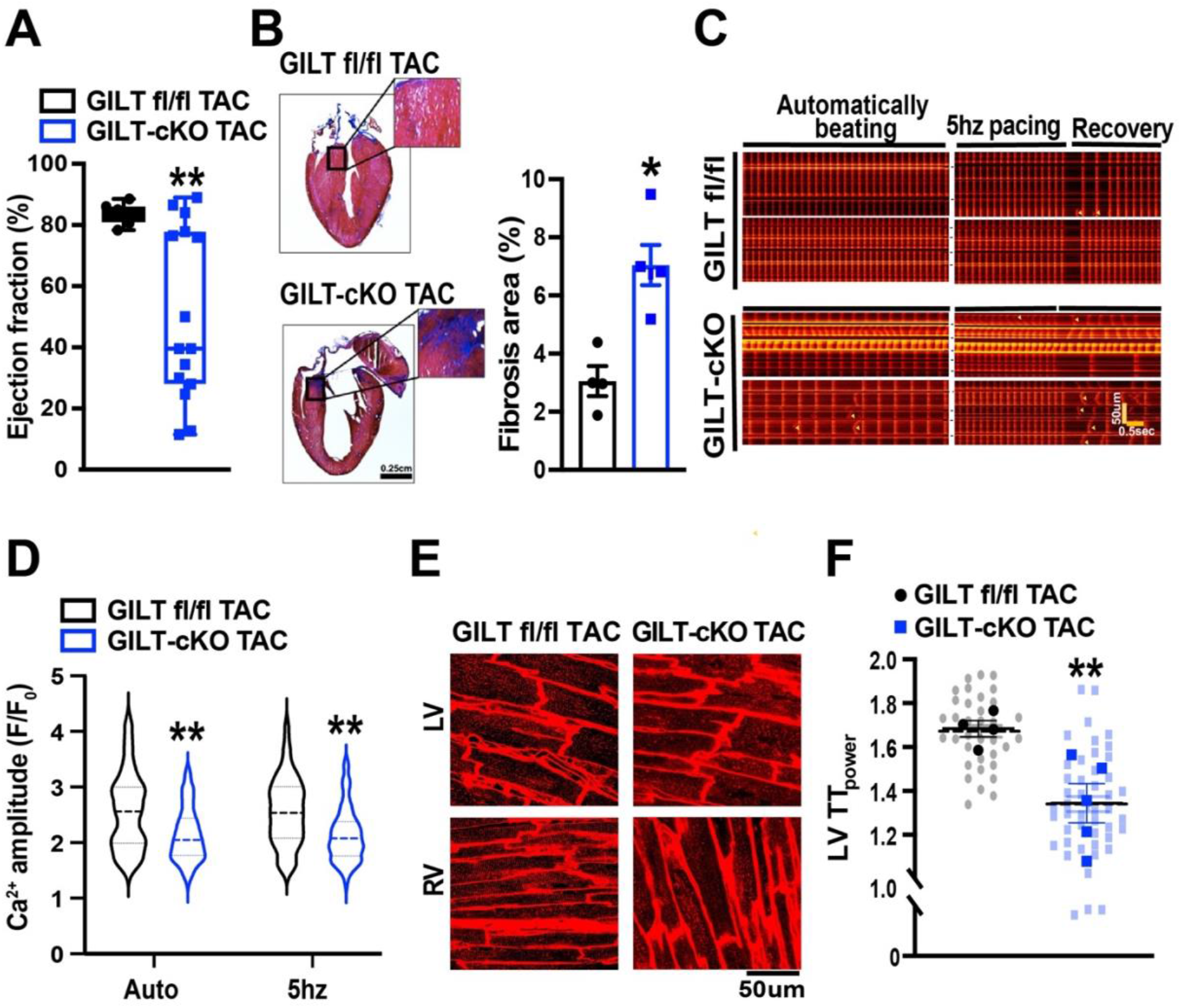
Cardiac GILT is necessary for maintaining structural and functional integrity of the heart. **A**. EF measured by echocardiography in GILT fl/fl and GILT-cKO male mice 5 weeks after TAC; n = 9-15 age-matched littermate mice per group. **B**. *Left*: Representative images of Masson’s trichrome staining in hearts from GILT fl/fl TAC and GILT-cKO TAC mice. *Right*: quantification of trichrome staining based on positive surface area normalized to total cardiac area; n= 4 age-matched littermate mice per group. **C**. Representative images of *in situ* confocal imaging of calcium in GILT fl/fl TAC and GILT-cKO TAC hearts perfused with Rhod-2AM prior to imaging; yellow triangles indicate abnormal calcium waves; scale bars: time = 0.5 sec, distance = 50 μm. **D**. Quantification of calcium amplitude in automatically beating (auto) and electrically stimulated (5 hz) hearts in (C); n=4 age-matched littermate mice per group. **E**. Representative images of *in situ* confocal imaging of T-tubules in GILT fl/fl TAC and GILT-cKO TAC hearts perfused with MM 4-64 prior to imaging; LV – left ventricle, RV – right ventricle; scale bar = 50 μm. **F**. Quantification of LV T-tubule power (TT_power_) in (E); n=4 age-matched littermate mice per group. Data are presented as means ± SEM. * Indicates genetic effect, as determined by Student’s t-test. * Indicates p-value < 0.05, ** indicates p-value < 0.01.

Obesity is associated with many cardiac intrinsic and extrinsic factors that can drive the progression of HF^8,9^. Therefore, we turned to a more specific cardiac stress model by utilizing transverse aortic constriction (TAC), a commonly used surgical procedure to induce pathological cardiac remodeling^51^. By 5 weeks post-TAC, GILT fl/fl male animals were able to sustain a compensated state (e.g., increased LV thickness albeit normal EF) potentially attributed to the C57BL/6J background, which is relatively resistant to cardiac stressors^52^ (Fig. 2A and SFig. 2D). In contrast, hearts from GILT-cKO mice became markedly dilated and contractile function was severely reduced, as evidenced by profound increases in end-diastolic/systolic volumes and reduced EF to ~ 40% (Fig. 2A and SFig. 2D). These findings were further supported by histological analysis, which showed that in GILT-cKO mice, TAC increased LV dilation and myocardial fibrosis as measured by Masson’s trichrome staining (Fig. 2B).

Defective Ca^2+^ handling is a hallmark of HF^53^. To investigate the role of GILT in cardiomyocyte calcium handling, a crucial mediator of normal heart contractility, we measured intracellular Ca^2+^ release in *in situ* epicardial myocytes from Langendorff-perfused intact TAC hearts using line-scan confocal imaging. To this end, Ca^2+^ images were acquired under spontaneous beating as well as under a fixed electric pacing rate (5 Hz). Although cardiomyocytes from GILT fl/fl hearts displayed nearly uniform autonomous beating, occasional Ca^2+^ waves were observed during recovery after 5-Hz electric pacing (Fig. 2C&D). In contrast, Ca^2+^ waves were more frequent and the amplitude of Ca^2+^ was impaired in GILT deficient hearts (Fig. 2C&D).

Transverse-tubules (T-tubules) integrate efficient electrical propagation and excitation-contraction coupling across ventricular myocytes^54^. To examine whether deletion of GILT affected the cardiomyocyte structural integrity, we carried out *in situ* imaging of T-tubules in intact hearts with TAC using confocal microscopy^55,56^. As shown in Fig. 2E&F, GILT-cKO hearts exhibited severe remodeling of T-tubules in both ventricles, a well-described signature of a failing heart^56^. Together, these data support the notion that cardiac GILT is crucial for normal heart function under physiological conditions and in response to pressure-overload stress.

### Loss of GILT in the heart disrupts cardiac metabolic reprogramming

To gain a broader understanding of GILT-dependent cardiac regulation, we performed an RNA-seq analysis using hearts from young (3 months of age) male GILT-cKO mice and their littermate controls. There was no major impact of cardiac GILT deletion on the lysosomal and autophagic transcriptional landscape (SFig. 3A&B). This finding is consistent with comparable activity of lysosomal enzymes, namely Hexosaminidase Subunit Beta (HexB) and Cathepsin B (CTSB) (SFig. 3C), measured under basal conditions. Pathway analysis further revealed immune response and metabolism as the top-enriched pathways that were induced and suppressed, respectively, in GILT-deficient hearts compared to GILT fl/fl hearts (Fig. 3A&B).

**Figure 3.**
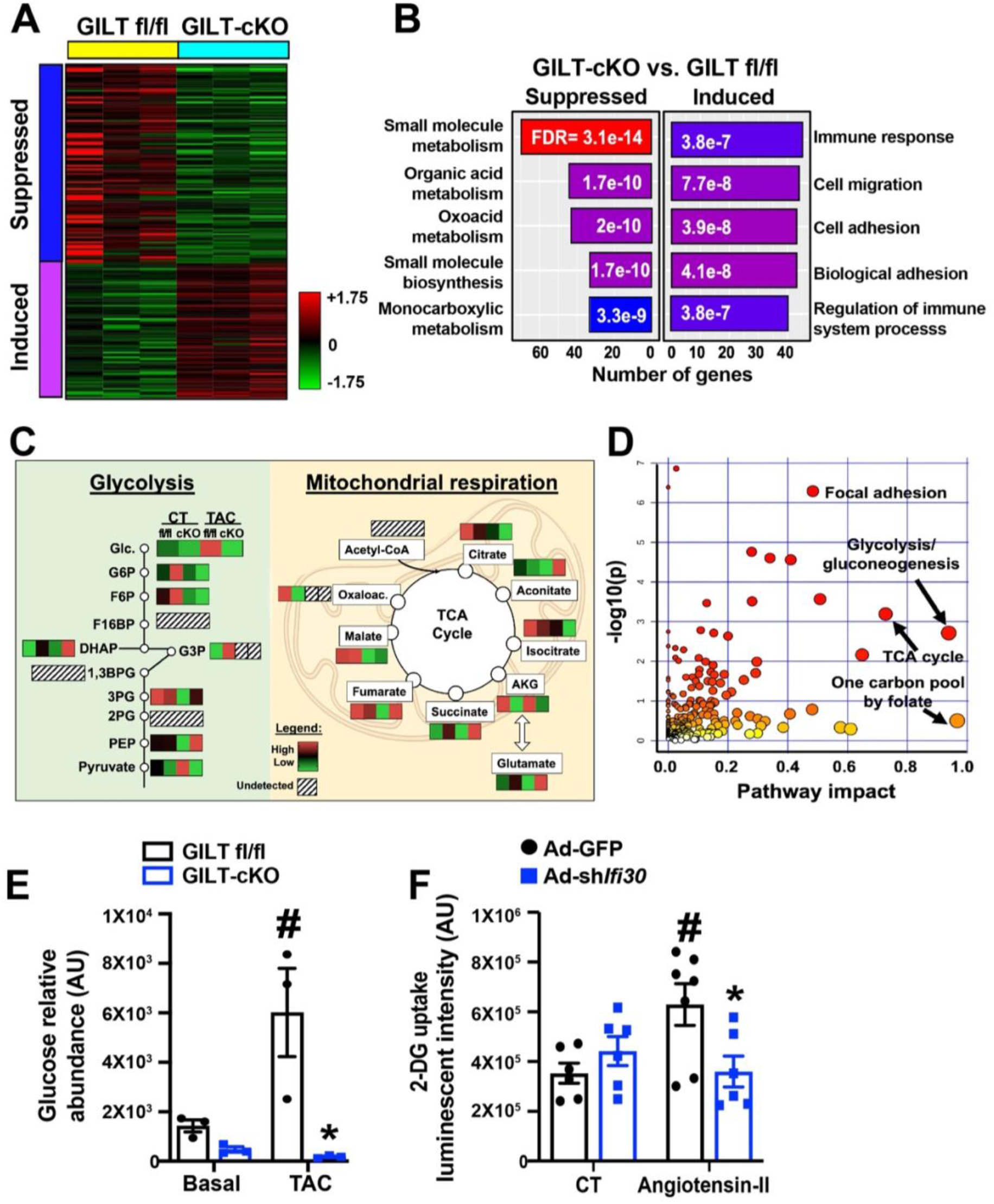
Deletion of cardiac GILT causes alterations in transcriptomic and metabolomic reprogramming in the heart. **A**. Bulk RNA-Seq analysis of hearts from GILT fl/fl and GILT-cKO mice at 3 months of age; differentially expressed transcripts were clustered into suppressed or induced functional groups; n=3 age-matched male mice per group; log_10_ fold change = +/− 1.75. **B**. Identification of top 5 significantly altered functional pathways in GILT fl/fl and GILT-cKO animals at 3 months of age; numbers within columns indicate false discovery rate (FDR); n = 3 age-matched male mice per group. **C**. Schematic summary of cardiac metabolomic analysis in GILT fl/fl and GILT-cKO under basal (CT) and TAC (5 weeks) conditions; n = 3 age-matched littermate male mice per group. **D**. Integration of transcriptomic and metabolomic signatures in GILT fl/fl and GILT-cKO mice under basal conditions analyzed by using Metaboanalyst online software. Highlighted pathways indicate top candidates with enriched both transcripts and metabolites that were significantly altered in GILT-cKO hearts compared to GILT fl/fl hearts; n=3 age-matched littermate male mice per group. **E**. Relative abundance of cardiac free glucose detected by GC-MS metabolomics in GILT fl/fl and GILT-cKO hearts under basal and TAC conditions. n = 3 age-matched littermate male mice per group. **F**. 2-DG uptake in HL-1 cells under basal (CT) conditions or in response to Angiotensin-II (1 uM); data were normalized to total protein. Data in (E&F) are presented as means ± SEM. *Indicates genetic effects and # indicates TAC effects in (E) and Angiotensin effects in (F). Statistical significance was determined by Two-way ANOVA followed by Tukey’s multiple comparisons test, p-value < 0.05.

Based on our transcriptomics findings, we first sought to determine the role of GILT in regulating cardiac metabolism under basal and TAC conditions. Inability to meet the high energy demands and altered patterns of substrate transport and utilization in the heart have been recognized as hallmarks of pathological metabolic remodeling in HF^6,9^. Therefore, to assess the impact of loss of GILT on cardiac metabolic homeostasis, we further carried out a steady-state cardiac metabolomic analysis using gas chromatography-mass spectrometry (GC-MS) in cardiac tissue. Deletion of cardiac GILT altered a wide array of metabolites in the heart under both basal (CT) and TAC (5 weeks) conditions (Fig. 3C&D). In general, there were significant alterations in the TCA cycle intermediates in GILT-cKO compared to GILT fl/fl animals under both basal and TAC conditions (Fig. 3C&D). Consistent with previous studies both in humans and mice^9,57^, in response to pressure overload cardiac levels of glucose and glycolytic metabolites were increased in GILT fl/fl mice (Fig. 3C&E). However, this response was diminished in GILT-cKO TAC hearts (Fig. 3C&E). To determine whether reduced glucose metabolites in GILT-deficient cardiac tissue was secondary to reduced glucose uptake, we determined glucose uptake analysis *in vitro* using an atrial cell line (Hl-1). Although suppression of *Ifi30* did not alter glucose uptake under basal conditions, it significantly decreased glucose uptake by cardiomyocytes in the presence of angiotensin-II challenge (Fig. 3F). We also observed a significant increase in levels of cardiac ketone bodies and amino acids in GILT-cKO TAC mice compared to GILT fl/fl mice in the setting of TAC (SFig. 3E&F). This finding is in line with a recent study reporting increased oxidation of ketone bodies and amino acids in failing human hearts^57,58^, implicating loss of GILT in the heart accelerates HF progression.

Reduction of mitochondrial ETC and elevation of mitochondrial ROS are hallmarks of HF-associated mitochondrial dysfunction^6^. Lysosomes control mitochondrial quality and thereby maintain mitochondrial function under both steady-state and stress conditions^19,59^. However, there was no reduction in protein levels within the mitochondrial oxidative phosphorylation (OxPhos) complex in GILT-cKO TAC mice compared to GILT fl/fl TAC mice (SFig. 4A&B). Increased mitochondrial reactive oxygen species (ROS) impair mitochondrial structure and function, which can contribute to cardiac dysfunction^60^. GILT-deficient primary cardiomyocytes exhibited increased levels of mitochondrial ROS both at baseline and in the presence of isoproterenol (Iso, a known inducer of mitochondrial ROS) relative to WT cells measured by using MitoSOX (Fig. 4A&B), a mitochondrial ROS probe^61^. To determine whether deletion of GILT altered cardiac mitochondrial respiratory capacity, we carried out high resolution respirometry analysis (Oxygraph-2k, Oroboros) in freshly dissected permeabilized cardiac fibers from unstressed GILT-cKO mice and littermate controls. Deletion of GILT diminished ADP-stimulated state-3 respiration in the presence of palmitoyl-carnitine and malate (Fig. 4C). This result is in line with our transcriptomic and metabolic findings showing that hearts from GILT-cKO animals had aberrant levels of TCA cycle metabolites as well as transcripts encoding metabolic regulators (Fig. 3B-D).

**Figure 4.**
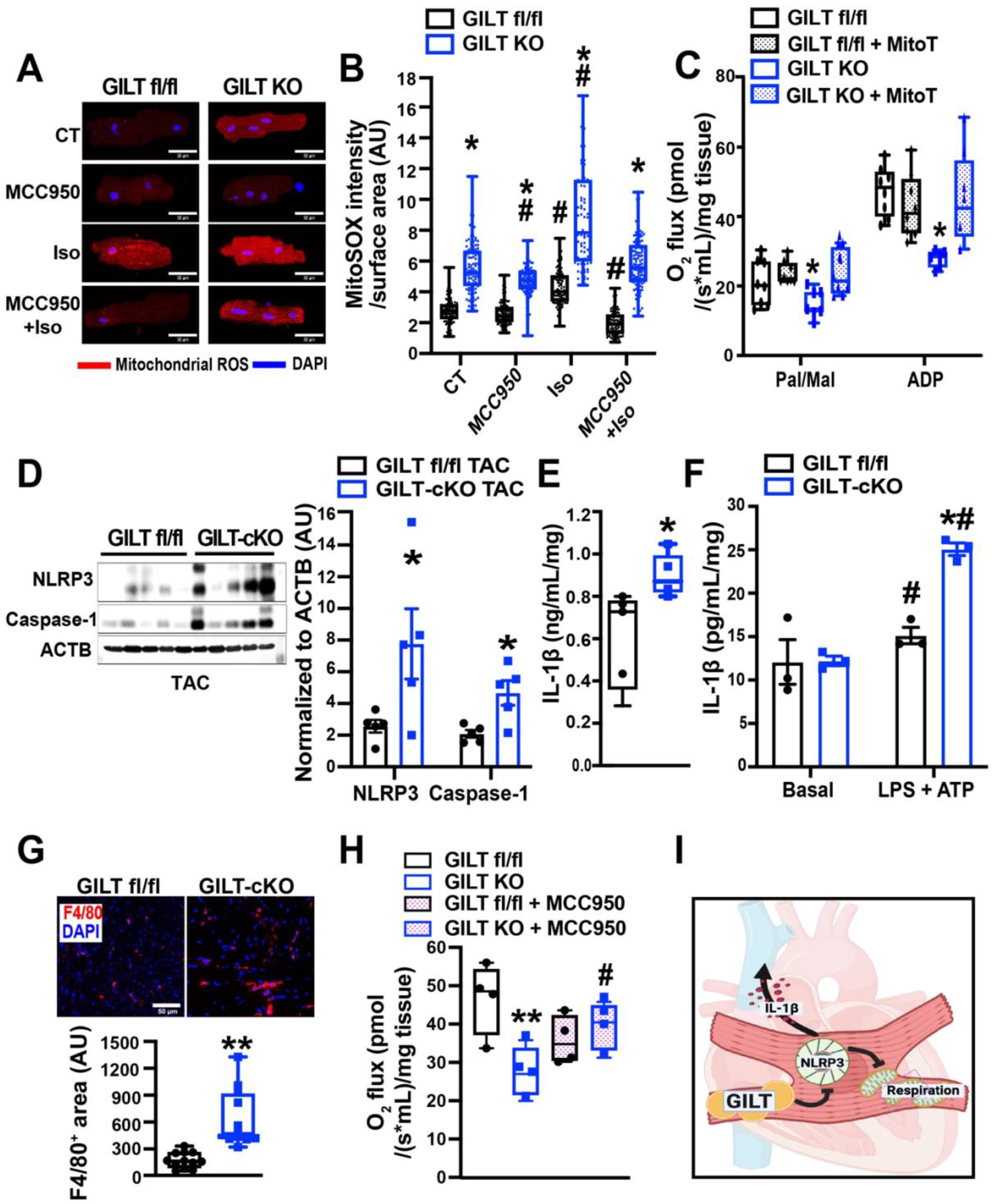
GILT-NLRP3 inflammasome axis modulates metabolic and inflammatory homeostasis in the heart. **A**. Representative confocal images of MitoSOX (red) and DAPI (blue) measured in primary cardiomyocytes from GILT fl/fl and GILT-cKO male mice. Cells were treated with MCC950 (1 μM) for 2 hours in the presence or absence of isoproterenol (Iso; 100 nM) for 15 min; scale bar = 50 μm. **B**. Quantification of MitoSOX intensity in (A); signal was normalized to cell surface area; n = 3 age-matched littermate male mice per group. **C**. Oxygen flux measured by high resolution respirometry (Oroboros) in permeabilized cardiac fibers isolated from GILT fl/fl and GILT-cKO mice; palmitoyl-carnitine (Pal; 0.075 mM), malate (Mal; 1 mM), and ADP (2.5 mM) were used as substrates. MitoTEMPO (MitoT, 50 μM) was added to the fibers for 1 hour prior to the analysis. The oxygen flux was normalized to wet weight of cardiac fibers; n = 3 age-matched littermate male mice per group. **D**. Western blots of and quantification of tested proteins in heart lysates from GILT fl/fl TAC and GILT-cKO TAC mice; n = 5 age-matched littermate male mice per group. **E**. ELISA of IL-1β measured in cardiac lysates from GILT fl/fl TAC and GILT-cKO TAC mice; n = 5 age-matched littermate male mice per group. **F**. ELISA of IL-1β measured in lysate from primary cardiomyocyte isolated from GILT fl/fl and GILT-cKO animals. The cells were treated with LPS (500 ng/mL) and ATP (5 mM); n = 3 age-matched littermate male mice per group. **G**. Representative images and quantification of immunohistochemistry analysis of F4/80-positive cells in the ventricular wall of the heart from GILT fl/fl TAC and GILT-cKO TAC mice; n = 4 age-matched littermate male mice per group; scale bar = 50 μm. **H**. Oxygen flux measured by high resolution respirometry (Oroboros) in permeabilized cardiac fibers isolated from GILT fl/fl and GILT-cKO mice; palmitoyl-carnitine (Pal; 0.075 mM), malate (Mal; 1 mM), and ADP (2.5 mM) were used as substrates. MCC950 (5 μM) was added 1 hour prior to the analysis **I**. Schematic model of GILT-dependent regulation of immuno-metabolic homeostasis in the heart. Data are presented as means ± SEM. * indicates genetic effects in cells/mice with same treatments; and # indicates treatment effects compared to the CT/Basal groups in cells/fibers with same genotype in (B and G). Statistic significances were determined by Student’s t-test in (E&G), and Two-way ANOVA followed by Tukey’s multiple comparisons test in (B, C, D F&H). * and # indicate p-value < 0.05, ** indicates p-value < 0.01.

We then addressed whether ameliorating mitochondrial ROS could protect against the GILT-deficiency-associated mitochondrial dysfunction. To this end, we treated cardiac fibers isolated from GILT fl/fl and GILT-cKO mice with Mito-TEMPO, a mitochondria-targeted antioxidant^62^, and then measured cardiac mitochondrial respiration. As shown Fig. 4C, treatment with Mito-TEMPO significantly ameliorated mitochondrial respiration in both GILT fl/fl and GILT-cKO cardiac fibers. Taken together, these data suggested that cardiac GILT-mediated inhibition of mitochondrial ROS protects mitochondrial function against cardiac stress.

### GILT shapes the inflammasome-mitochondria axis in the context of HF

Our transcriptomic analysis revealed that loss of GILT promotes a pro-inflammatory signature in the heart under basal conditions (Fig. 3B). The NLR family pyrin domain containing 3 (NLRP3) inflammasome is a key element in the cardiac inflammatory response to a cardiac injury^63–65^. Notably, the lysosome plays a key role in both the priming and assembly phases of the inflammasome^66–68^. Indeed, we found that loss of GILT in cardiomyocytes increased TAC-induced NLRP3 and Caspase-1 expression (Fig. 4D&E). Similar results were observed in mice challenged with a HFD (SFig. 4C). Sustained activation of proinflammatory signaling cascades and release of IL-1β results in the recruitment and activation of inflammatory immune cells, another critical factor in the progression of HF^69^. Given that inflammasome activation is a primary source of IL-1β^69^, we measured IL-1β production in heart lysates from GILT-cKO TAC mice and their GILT fl/fl TAC controls. As shown in Fig. 4F, the level of IL-1β was significantly increased in GILT-cKO hearts after TAC. Next, we asked whether this effect was in part cell autonomous. To this end, GILT-deficient primary cardiomyocytes were exposed to lipopolysaccharide (LPS), followed by a short exposure to ATP, a commonly used paradigm to activate the inflammasome^70^. We found that GILT deficiency elevates cardiomyocyte IL-1β secretion, indicating a cell-autonomous effect of GILT deletion on inflammasome activity (Fig. 4G). Together, these results indicate an anti-inflammatory role of GILT in HF progression, which is mediated in part by regulating the NLRP3-IL-1β axis^69^.

Reciprocal modulation of mitochondrial dysfunction and NLRP3 inflammasome activation has been evident in diverse forms of cardiac pathologies^64,65^. To dissect the causality of the GILT-inflammasome axis and GILT-mitochondrial axis, we treated primary cardiomyocytes with a potent NLRP3 inhibitor, MCC950, which blocks the release of IL‑1β induced by NLRP3 activators^71^, and measured its effect on mitochondrial ROS production. Notably, inhibiting NLRP3 reduced both basal and isoproterenol-induced mitochondrial ROS in GILT-deficient cardiomyocytes (Fig. 4A&B), indicating that NLRP3 activation is an upstream process that induces GILT deficiency-associated mitochondrial respiratory defects. In line with the elevation of IL‑1β production, immunohistochemical (Fig. 4G) and flow cytometry analyses (SFig. 4D) further revealed that GILT deficiency significantly increased macrophage infiltration in the heart after TAC. To further assess the causality of the NLRP3-driven mitochondrial defective respiration in GILT-cKO mice, we treated freshly isolated permeabilized cardiac fibers from WT and GILT KO mice with MCC950 annd measured mitochondrial respiration. As shown in Fig. 4H, inhibition of NLRP3 partially rescues the defective mitochondrial respiration in GILT-deficient cardiac fibers. Together, these data demonstrate that cardiac GILT controls the balance between metabolic and inflammatory remodeling in HF in part through modulating NLRP3 inflammasome action (Fig. 4I).

## DISSCUSSION

Heart failure progression is associated with, and in part driven by, a myriad of cardiac structural (hypertrophic growth) and functional (metabolic and immune) pathological changes within the heart. Understanding the interplay between these changes in cardiomyocytes in response to an injury may reveal new therapeutic avenues for HF patients. Here, we demonstrate a previously unrecognized role of lysosomal function in linking cardiac inflammation with metabolic remodeling in the setting of pressure-overload-induced HF.

Lysosomal degradation and autophagy are essential for cellular quality control in the heart in health and disease^34,35,72,73^. In the context of cardiac pathologies, such as ischemia/reperfusion (I/R) and myocardial infarction (MI), activation of autophagy can be beneficial or detrimental, depending on the environmental context^35,36,39,40,74,75^. Under basal conditions, we observed no significant alterations in cardiac systolic function in mice with GILT deletion. However, loss of GILT in the heart exacerbated cardiac dysfunction in aged, obese and pressure-overloaded hearts. We hypothesize that GILT-mediated lysosome function plays important roles in maintaining structural and functional integrity of the heart, which delays an irreversible injury after prolonged exposure to stress (i.e., TAC or HFD). Further studies will be necessary to evaluate spatio-temporal regulation of GILT-dependent cardiac remodeling in response to stress.

Mounting evidence demonstrates that dysregulation of the immune response contributes to the pathogenesis of HF and its associated adverse clinical outcomes^5,13^. The inflammasome is a protein complex that senses injury/damage signals and triggers an inflammatory response ^76^. Nod-like receptor protein (NLRP)3 inflammasomes is a widely characterized inflammasome sensor in the heart. Of note, inhibition of NLRP3 activation and its associated activation of IL-1β production have been shown to improve cardiac pathologies in both experimental and clinical studies^64,65^.

There are two steps involved in NLPR3 activation: the priming step, which induces the expression of inflammasome components and the activation/triggering step of NLPR3, in which the components assemble and propagate downstream actions^76,77^. We found that cardiac GILT deletion exacerbated TAC-induced expression of Caspase-1 and IL-1β production in the heart. However, we did not observe induction of *Nlrp3* and the associated components in our transcriptomic analysis (SFig. 3D), indicating that the increased inflammasome activation in GILT-cKO hearts may not occur at the priming stage. Accumulating evidence demonstrates that mitochondrial dysfunction/ROS overproduction is potent cellular cue for triggering NLRP3 inflammasome activation^68^. In our study, we found that chemical inhibition of NLRP3 activation reduced mitochondrial ROS production in primary cardiomyocytes, indicating that GILT-mediated NLRP3 action could be an upstream regulator of the of mitochondrial ROS signaling cascade, thus we postulated this might be in part due to IL-1β overproduction-associated mitochondrial dysfunction.

Lysosomal membrane damage triggers K^+^ efflux and leaking of lysosomal contents, which activate NLRP3^68,77,78^. In the heart, lysosomal stability is significantly altered at the site of injured myocardium within 1 hour of ischemia^79^; further, patients with MI also show compromised lysosomal stability^80^. Although it is appreciated that perturbations of lysosomes lead to NLRP3 activation in non-cardiac cells, such as macrophages^67^, fewer studies have explored the cardiac lysosome-inflammasome axis in the context of HF. In addition to enzymatic action, the lysosome-autophagy machinery also modulates non-conventional IL-1β secretion^81^ and inflammasome activation in immune cells (i.e. macrophages and neutrophils)^39,82,83^. The lysosome also serves as an intracellular platform for metabolic and calcium homeostasis^19,84^. In fact, recent studies showed that lysosome-mediated calcium mobilization activates the NLRP3 inflammasome^67,85^. Moreover, disruption of cardiomyocyte calcium flux and activation of CaMKII directly activates NLRP3 in atrial fibrillation^63,86^. Future experiments will elucidate which of these signaling cascades directly link the lysosome-NLRP3 inflammasome axis in cardiomyocytes.

Progression of HF involves profound cardiac metabolic remodeling characterized by decreased energy production, altered fuel/substrate utilization, and disrupted cellular repair processes^8,9^. The contractile and metabolic functions of the heart require massive mitochondrial energy production; accordingly, mitochondrial dysfunction is a hallmark of HF. Clinically, improving mitochondrial function in cardiomyocytes has been tested as a potential approach to improve the function of a failing heart^6,87,88^. Lysosomes play a critical role in mitochondrial quality control and substrate supply ^19^. We observed that cardiac GILT deletion impaired mitochondrial respiration without altering the OxPhos protein complex (SFig. 3A&B). Although to this point we cannot rule out a possibility that defective oxidative phosphorylation is due to altered mitochondrial ultrastructure in the Gilt-deficient heart, our data reveal the concurrence of excessive mitochondrial ROS and mitochondrial dysfunction in the setting of HF. Mitophagy plays essential roles in shaping mitochondrial respiration^69^. However, transcripts of the genes involved in macroautophagy and mitophagy were not significantly altered in GILT cKO hearts under basal conditions (SFig. 3A&B). Instead, we observed an evident effect of deleting GILT on the immuno-metabolic profile (Fig. 3B&C). Therefore, we postulate that loss of GILT-mediated lysosomal function perturbs immuno-metabolic pathways in the cardiomyocyte and renders the heart susceptible to pressure overload-induced injury. Indeed, recent studies from Kenny et al.^40^ and Trivedi et al.^89^, which utilized gain-of and loss-of cardiac TFEB function, respectively, all point to an autophagy-independent role of metabolic regulation by the lysosome.

Under normal conditions, fatty acid oxidation (FAO) serves as the primary energy source for ATP production in cardiomyocytes *in vivo*. In contrast, during HF, cardiomyocytes shift from FAO toward glycolytic metabolism that is uncoupled from pyruvate oxidation^9,90–93^. Our metabolomic analysis revealed that GILT deficiency significantly lowered levels of free glucose and glycolytic intermediates in the heart, indicating altered metabolic reprogramming in the knockout animals, particularly in response to hemodynamic stress. The lysosome plays an important role in the regulation of glucose and lipid transporters, such as glucose transporter type 4 (GLUT4) and cluster of differentiation 36 (CD36), respectively in the adipocyte and hepatocyte^66,94–96^. The lysosome is also the subcellular site of glycogenolysis^97^. Ultimately, the importance and temporal roles of lysosomes in modulating these key metabolic features of the cardiomyocyte warrant further investigation.

Proper immune responses and metabolic reprogramming are essential for the heart under conditions of stress. Our study demonstrates a previously unrecognized role of GILT in regulating the immuno-metabolic interplay in the heart under healthy and stress conditions. These findings offer a platform for dissecting the contribution of lysosomal dysfunction in HF and identify potential targets for therapeutic intervention for patients with HF.

## Supporting information

Main text-2023

## ARTICLE INFORMATION

## Acknowledgments

We thank Dr. Karen Hastings at the University of Arizona and Peter Cresswell at the Yale School of Medicine for providing the GILT^−/−^ mouse. We thank the University of Iowa Cardiology Mouse Phenotyping Core Laboratory for performing transthoracic echocardiography as well as Metabolic Core Facility and Iowa Institute of Human Genetics at the University of Iowa for performing metabolomics and RNA-Seq analyses respectively. We also thank Dr. Vitor Lira at University of Iowa for technical support and Dr. Sara C. Sebag for editing this manuscript, and all Yang laboratory members for supporting this project.

## Author contributions

M.L. and L.Y. designed the study and wrote the manuscript. M.L., J. W., Q.Q., B.C., Z.Z., E.B., Y.Z., and D.D.H. performed the experiments. E.D.A. and L.S.S. provided critical reagents and scientific suggestions on the manuscript. L.Y. conceived and supervised the study.

---

**SFig. 1.**
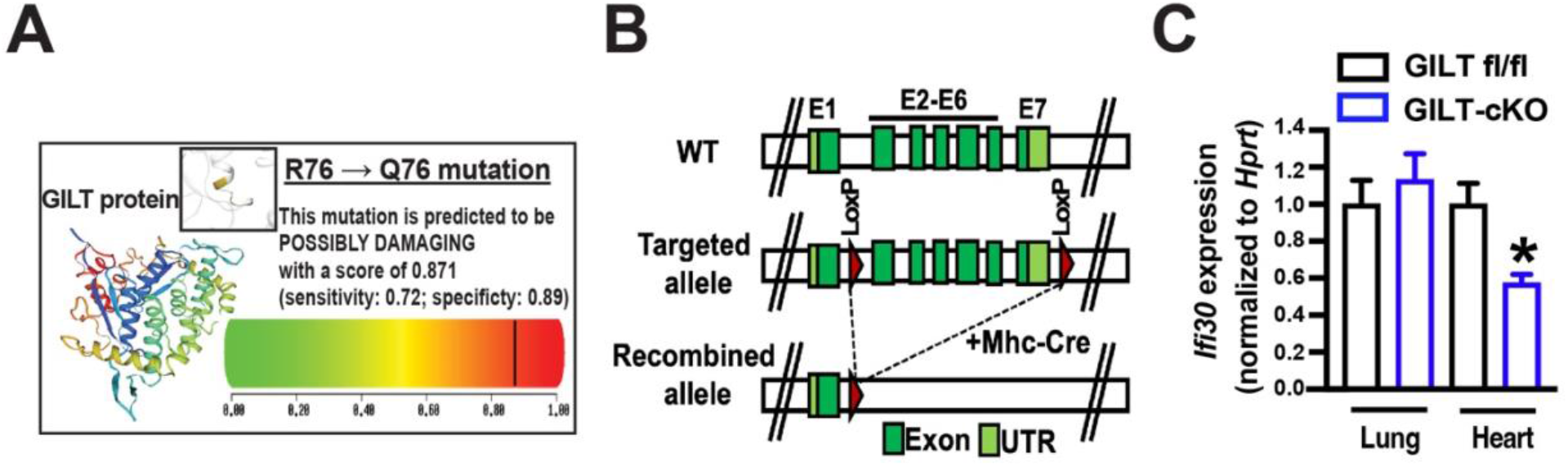
Cardiac GILT deletion mouse model. **A**. *in silico* analysis ^47^ for deleterious effects of the SNP on GILT’s enzymatic activity. **B**. Schematic diagram of the strategy to delete *Ifi30* in cardiomyocytes. **C**. Expression of the *Ifi30* transcript measured by qRT-PCR from lung and heart of GILT fl/fl mice and mice with cardiac deletion of GILT (GILT-cKO); n = 3 aged-matched GILT -cKO and littermate control (GILT fl/fl) mice. Data are presented as means ± SEM in (C), and statistic significances were determined by Student’s t-test, * indicate p-value < 0.05.

**SFig. 2.**
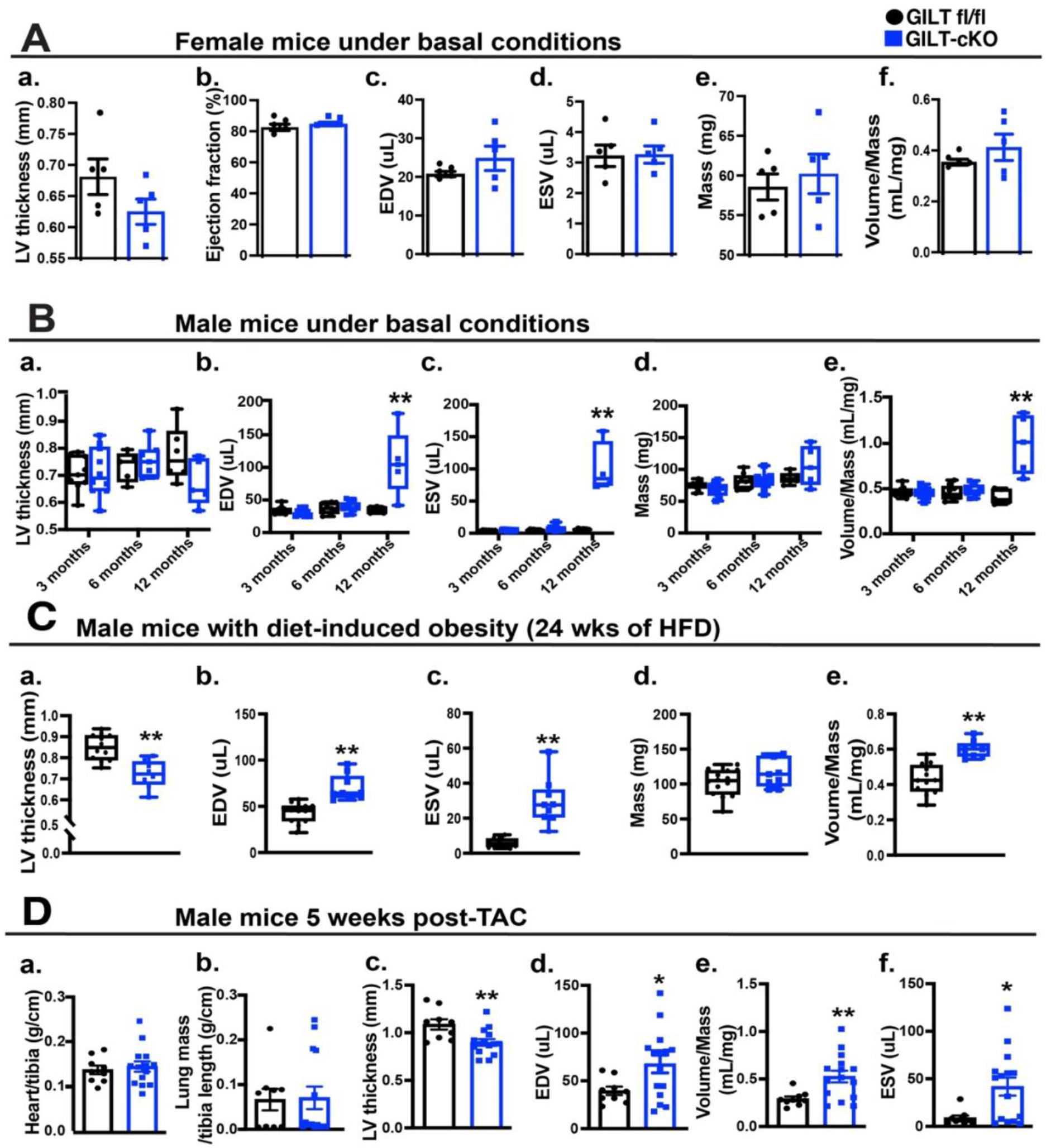
Cardiac GILT deletion leads to heart dysfunction. Echocardiographic parameters measured in GILT fl/fl or GILT-cKO female mice (3 months old) **(A)**, male mice at different ages **(B)**, male mice on a HFD (**C;** 24 weeks on diet), and mice subjected to TAC **(D)**. Data are presented as means ± SEM. * indicates genetic effects as determined by Student’s t-test. * Indicates p-value < 0.05, ** indicates p-value < 0.01.

**SFig. 3.**
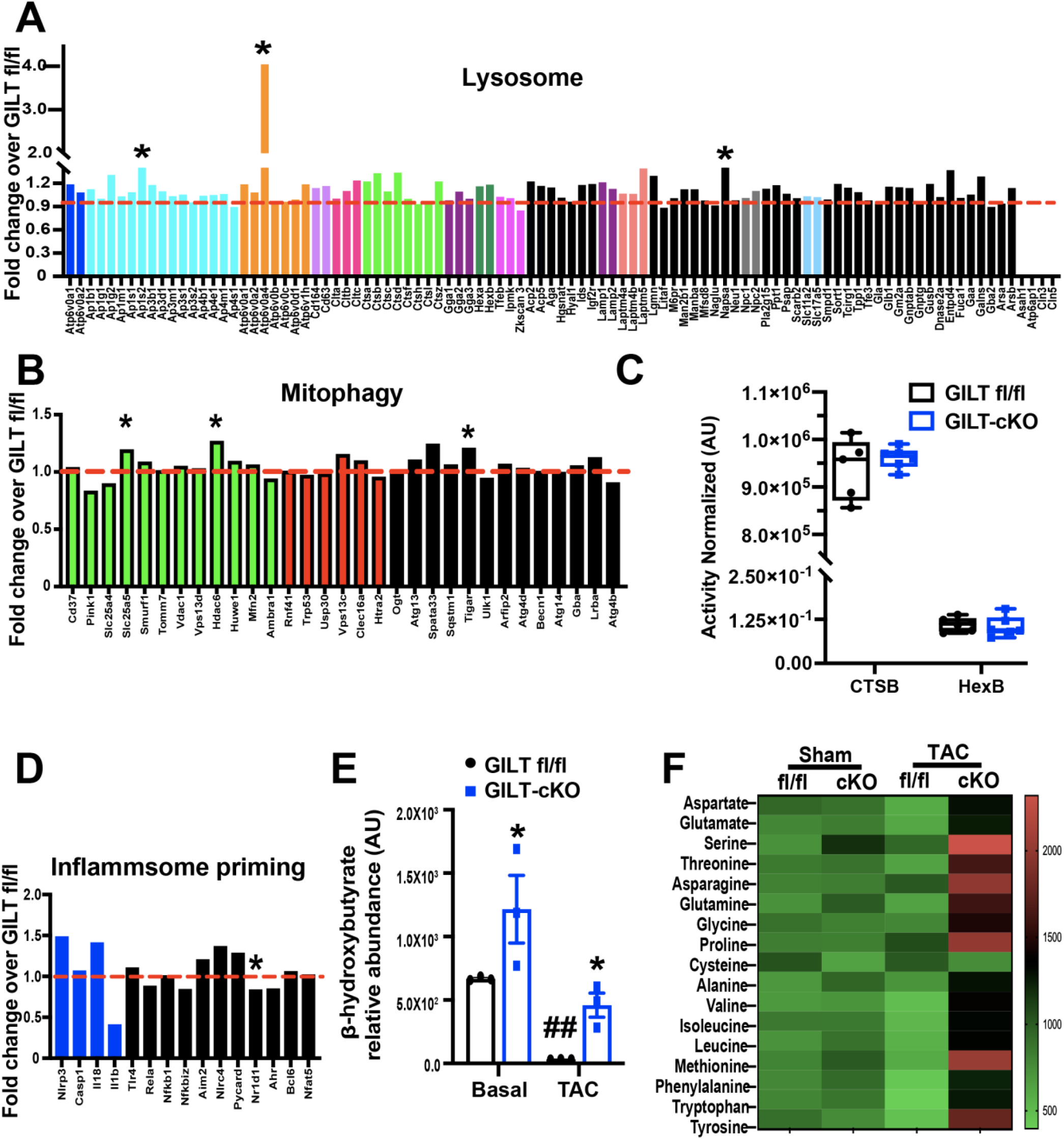
The effects of loss of cardiac GILT on cardiac transcriptome and metabolome. Lysosomal/ autophagic transcriptome (**A**), mitophagic transcriptome (**B**) and inflammasome signature (**D**) in hearts from GILT fl/fl and GILT-cKO mice at 3 months of age. **C**. CTSB and HexB enzyme activity measured in GILT fl/fl and GILT-cKO hearts under basal conditions. n = 3-6 age-matched littermate male mice per group. **D**. Transcripts associated with inflammasome priming analyzed by RNA-Seq in hearts from GILT fl/fl and GILT-cKO mice at 3 months of age. **E&F**. Relative abundance of cardiac ketone and amino acids in GILT fl/fl and GILT-cKO hearts under basal and TAC conditions. Data are presented as means ± SEM. * indicates genetic effects in mice with same treatments; and # indicates treatment effects compared in mice with the same genotype. Statistical significance was determined by Two-way ANOVA followed by Tukey’s multiple comparisons test. *Indicates p-value < 0.05, ** and ## indicates p-value < 0.01.

**SFig. 4.**
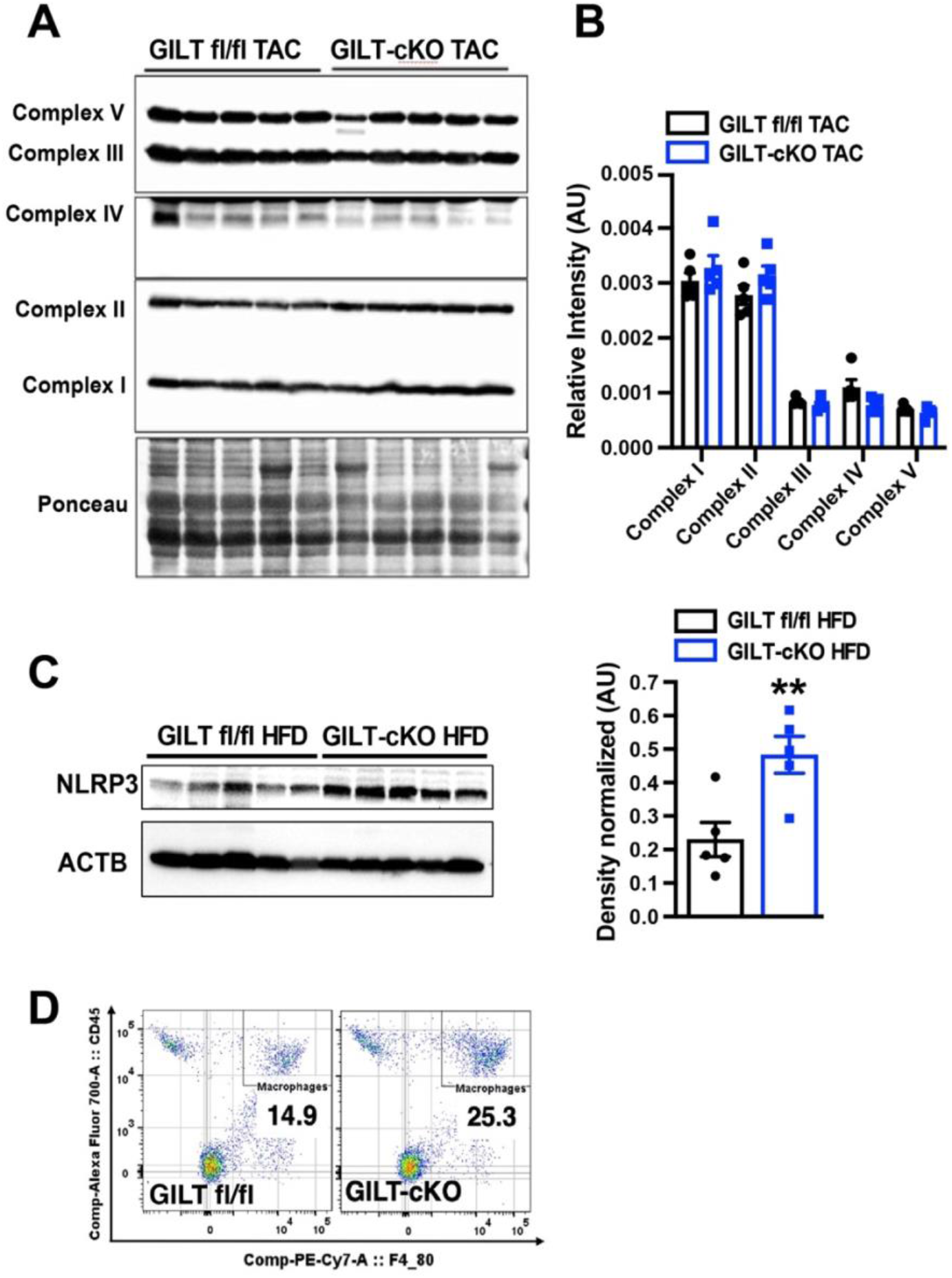
Effects of cardiac GILT deletion on immuno-metabolic profile in the heart. **A&B**. Representative western blots and quantification of mitochondrial oxidative phosphorylation (OxPhos) complexes in GILT-cKO TAC mice compared to GILT fl/fl TAC (5 weeks) mice at 15 weeks of age. n = 5 male littermate male mice per group. **C**. Western blots and quantification of NLPR3 in GILT-cKO AND GILT fl/fl on a HFD (24 weeks on diet). n = 5 male mice/group. **D**. Flow cytometry analysis of single cells isolated from 15-week-old GILT fl/fl and GILT-cKO hearts; quantification represents a gated population of CD45+/F4/80+ cells (macrophages). Data are presented as means ± SEM. *Indicates genetic effects as determined by Student’s t-test. Two-way ANOVA followed by Tukey’s multiple comparisons test was used for (B). *Indicates p-value < 0.05, ** indicates p-value < 0.01.

