## Supplementary material for "Gamma-interferon-inducible lysosomal thiol reductase maintains cardiac immuno-metabolic homeostasis in heart failure": Main text-2023

### Supplemental Figure 1

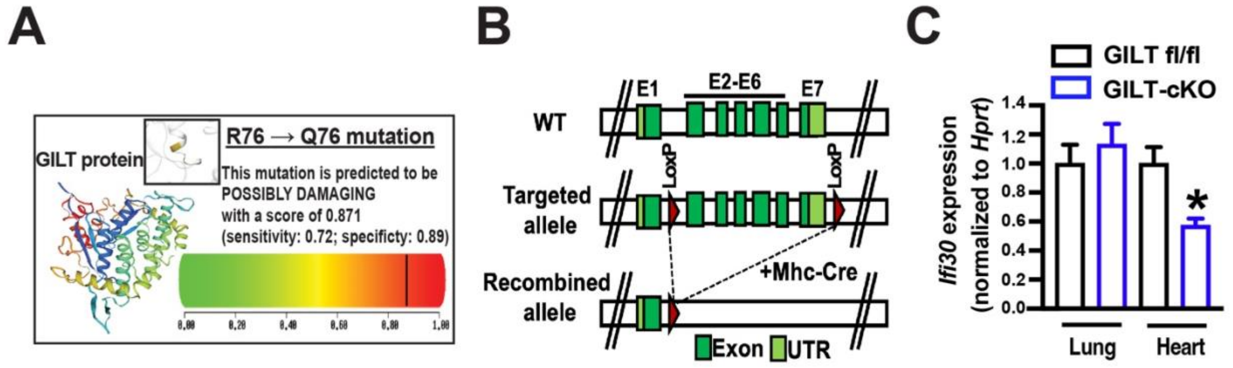


**SFig. 1. Cardiac GILT deletion mouse model. A**. *in silico* analysis [47] for deleterious effects of the SNP on GILT's enzymatic activity. **B.** Schematic diagram of the strategy to delete *Ifi30* in cardiomyocytes. **C.** Expression of the *Ifi30* transcript measured by qRT-PCR from lung and heart of GILT fl/fl mice and mice with cardiac deletion of GILT (GILT-cKO); n = 3 aged-matched GILT -cKO and littermate control (GILT fl/fl) mice. Data are presented as means ± SEM in (C), and statistic significances were determined by Student's t-test, * indicate p-value < 0.05.

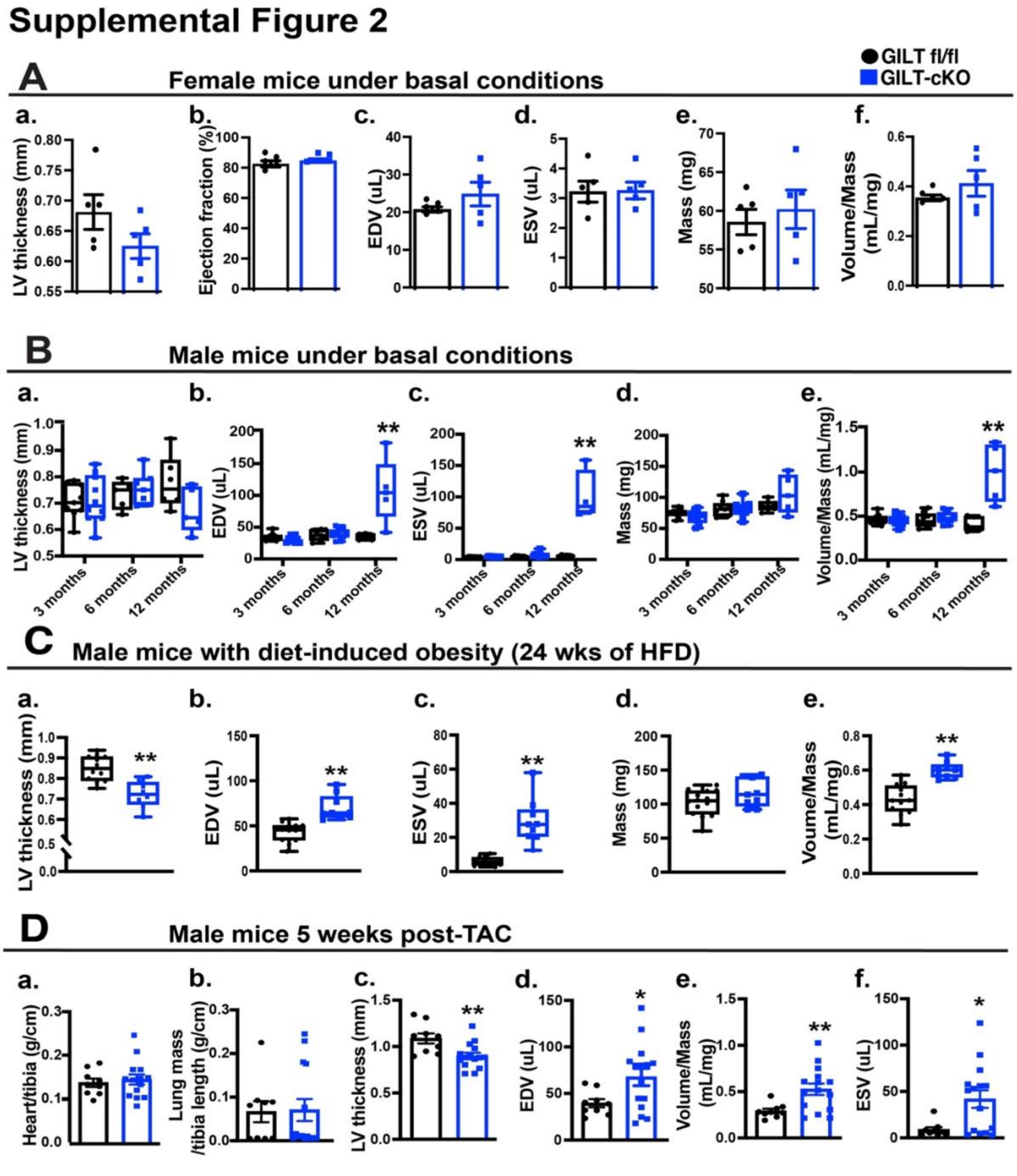


**SFig. 2. Cardiac GILT deletion leads to heart dysfunction.** Echocardiographic parameters measured in GILT fl/fl or GILT-cKO female mice (3 months old) **(A),** male mice at different ages **(B)**, male mice on a HFD (**C;** 24 weeks on diet), and mice subjected to TAC **(D)**. Data are presented as means ± SEM. * indicates genetic effects as determined by Student's t-test. * Indicates p-value < 0.05, ** indicates p-value < 0.01.

### Supplemental Figure 3

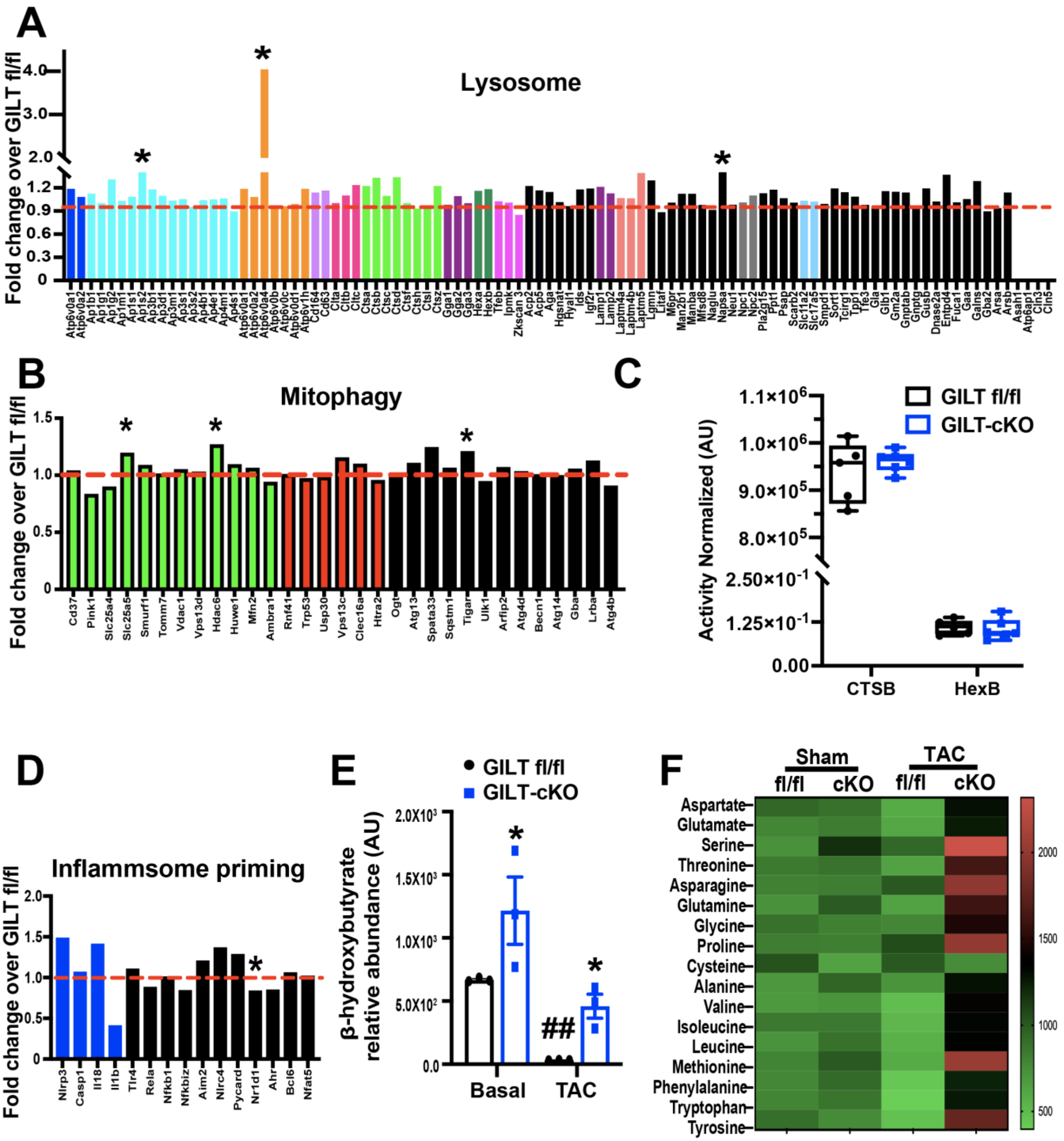


**SFig. 3. The effects of loss of cardiac GILT on cardiac transcriptome and metabolome.** Lysosomal/ autophagic transcriptome (**A**), mitophagic transcriptome (**B**) and inflammasome signature (**D**) in hearts from GILT fl/fl and GILT-cKO mice at 3 months of age. **C**. CTSB and HexB enzyme activity measured in GILT fl/fl and GILT-cKO hearts under basal conditions. n = 3-6 age-matched littermate male mice per group. **D.** Transcripts associated with inflammasome priming analyzed by RNA-Seq in hearts from GILT

fl/fl and GILT-cKO mice at 3 months of age. **E&F**. Relative abundance of cardiac ketone and amino acids in GILT fl/fl and GILT-cKO hearts under basal and TAC conditions. Data are presented as means ± SEM. * indicates genetic effects in mice with same treatments; and # indicates treatment effects compared in mice with the same genotype. Statistical significance was determined by Two-way ANOVA followed by Tukey's multiple comparisons test. *Indicates p-value < 0.05, ** and ## indicates p-value < 0.01.

#### Supplemental Figure 4

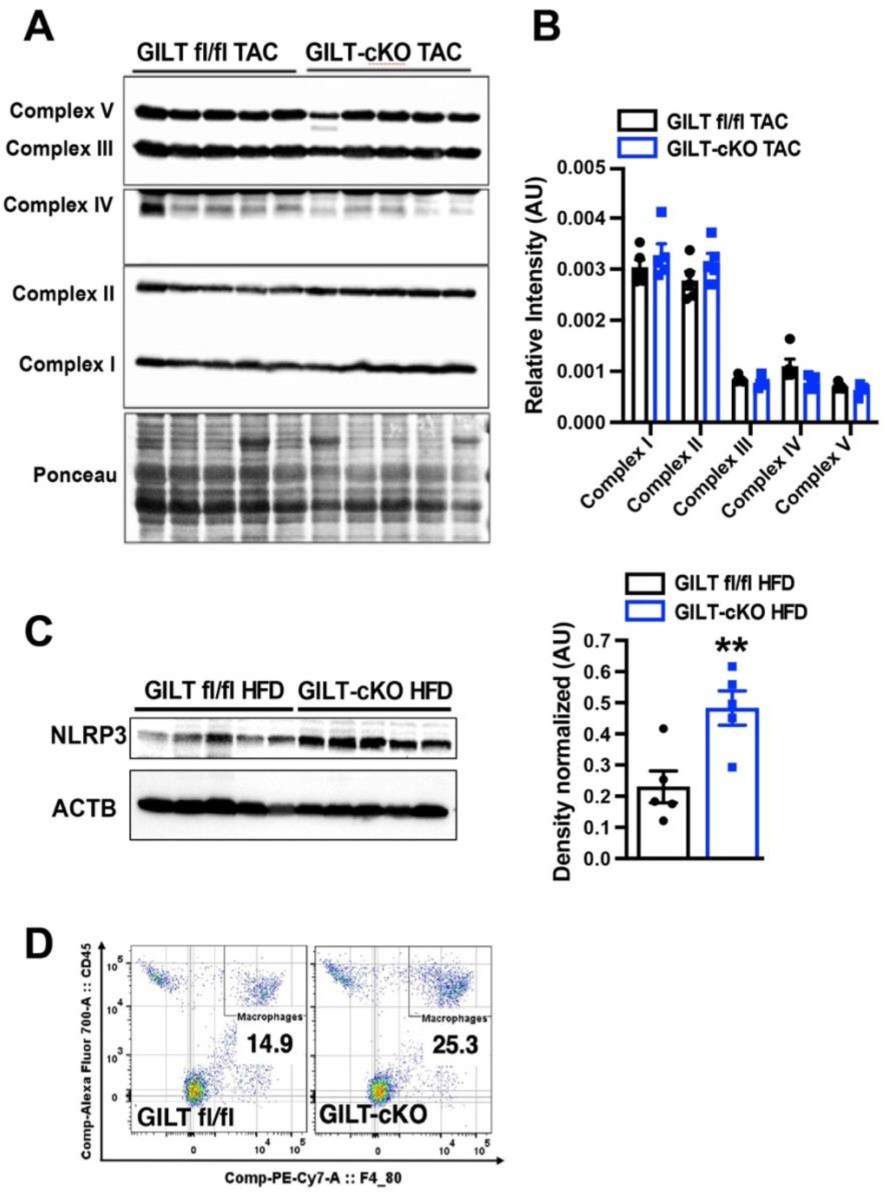


**SFig. 4. Effects of cardiac GILT deletion on immuno-metabolic profile in the heart. A&B.** Representative western blots and quantification of mitochondrial oxidative phosphorylation (OxPhos) complexes in GILT-cKO TAC mice compared to GILT fl/fl TAC (5 weeks) mice at 15 weeks of age. n = 5 male littermate male mice per group. **C.** Western blots and quantification of NLPR3 in GILT-cKO AND GILT fl/fl on a HFD (24 weeks on diet). n = 5 male mice/group. **D.** Flow cytometry analysis of single cells isolated from 15-week-old GILT fl/fl and GILT-cKO hearts; quantification represents a gated population of CD45+/F4/80+ cells (macrophages). Data are presented as means ± SEM. *Indicates genetic effects as determined by Student's t-test. Two-way ANOVA followed by Tukey's multiple comparisons test was used for (B). *Indicates p-value < 0.05, ** indicates p-value < 0.01.
